# Age-related neurodegeneration and cognitive impairments of NRMT1 knockout mice are preceded by misregulation of RB and expansion of the neural stem cell population

**DOI:** 10.1101/2021.03.15.435479

**Authors:** James Catlin, Leandro N. Marziali, Benjamin Rein, Zhen Yan, M. Laura Feltri, Christine E. Schaner Tooley

## Abstract

N-terminal methylation is an important post-translational modification that regulates protein/DNA interactions and plays a role in many cellular processes, including DNA damage repair, mitosis, and transcriptional regulation. Our generation of a constitutive knockout mouse for the N-terminal methyltransferase NRMT1, demonstrated its loss results in severe developmental abnormalities and premature aging. As premature aging is often accompanied by neurodegeneration, we more specifically examined how NRMT1 loss affects neural pathology and cognitive behaviors. Here we find that *Nrmt1*^-/-^ mice exhibit postnatal enlargement of the lateral ventricles, age-dependent striatal and hippocampal neurodegeneration, memory impairments, and hyperactivity. These morphological and behavior abnormalities are preceded by alterations in neural stem cell (NSC) development. Depletion of quiescent NSC pools in *Nrmt1*^-/-^ mice is concurrent with expansion of intermediate progenitor and neuroblast pools. These phenotypes are similar to those seen with loss of the NRMT1 target retinoblastoma protein (RB), and we see that NRMT1 loss leads to derepression of RB target genes and abnormal RB phosphorylation and degradation. As also seen with RB loss, neurons in *Nrmt1*^-/-^ mice fail to exit cell cycle and ultimately undergo NOXA-mediated apoptosis, indicating that early misregulation of RB in *Nrmt1*^-/-^ mice promotes premature NSC proliferation and contributes to subsequent neurodegenerative phenotypes.

## Introduction

N-terminal methylation of the alpha amino group (Nα-methylation) is a highly conserved post-translational modification (PTM) that regulates protein stability and protein/DNA interactions (Pettigrew and Smith 1977; Chen et al. 2007; Faughn et al. 2018). Nα-methylation has been shown to protect against digestion by cellular proteases (Pettigrew and Smith 1977), and Nα- trimethylated proteins receive a pH-insensitive, net positive charge that is thought to help facilitate electrostatic binding with negatively charged phosphate groups in the DNA backbone (Stock et al. 1987; Chen et al. 2007; Dai et al. 2013; Cai et al. 2014). Though this modification was discovered more than 40 years ago, and is the second most common Nα-PTM, we are just beginning to discover the many different ways it impacts normal development and disease.

Ten years ago, we identified the first mammalian Nα-methyltransferase, NRMT1 (<u>N</u>-terminal <u>R</u>CC1 <u>m</u>ethyltransferase 1) (Tooley et al. 2010). NRMT1 is a highly conserved enzyme that N-terminally trimethylates proteins based on a characterized Nα-consensus sequence (Tooley et al. 2010; Petkowski et al. 2012). This consensus sequence predicts more than 300 possible substrates for NRMT1 (Petkowski et al. 2012). Verified targets play roles in mitosis (RCC1), cell cycle (RB), DNA damage repair (DDB2), chromatin organization (CENP-B, CENP-A), cell motility (MYL9), and transcriptional regulation (SET) (Tooley et al. 2010; Petkowski et al. 2012; Bailey et al. 2013; Dai et al. 2013; Cai et al. 2014). Accordingly, loss of NRMT1 results in a variety of different cellular phenotypes, including multi-polar spindles, aneuploidy, impaired DNA damage repair, and altered cell growth (Chen et al. 2007; Tooley et al. 2010; Cai et al. 2014; Bonsignore et al. 2015a). In breast cancer cells, NRMT1 acts as a tumor suppressor, and its loss increases cell proliferation, cell migration, colony formation, and xenograft tumor development (Bonsignore et al. 2015a). Our recent creation of the first NRMT1 knockout (*Nrmt1*^-/-^) mouse has also revealed its importance in development and aging. Approximately 40% of *Nrmt1*^-/-^ mice die within the first month, and of those that survive, only 30% live past 6 months (Bonsignore et al. 2015b). Those that survive past 6 months exhibit phenotypes associated with premature aging, including early graying, kyphosis, dermal fibrosis, and sensitivity to DNA damaging agents (Bonsignore et al. 2015b).

As DNA damage accumulation and premature aging phenotypes often precede or are accompanied by neurodegeneration (Coppede and Migliore 2010), we wanted to look more specifically at how NRMT1 loss affects age-related neural pathology and cognitive behaviors. Here, we show the lateral ventricles of *Nrmt1*^-/-^ mice enlarge between 14 days and 6 weeks of age and this is accompanied by a significant reduction in volume of the adjacent striatum. We also see significant degeneration of hippocampal neurons in the third *Cornu Ammonis* area (CA3) and dentate gyrus (DG) starting at 3 months old. Corresponding to these morphological differences in the striatum and hippocampus, behavioral analyses showed *Nrmt1*^-/-^ mice exhibit hyperactivity and impaired short- and long-term memory, respectively.

The subventricular zone (SVZ) of the lateral ventricles and the subgranular zone (SGZ) of the dentate gyrus in the hippocampus contain the postnatal stem cell niches (Obernier and Alvarez-Buylla 2019). As these regions both show significant morphological abnormalities in *Nrmt1*^-/-^ mice, we next examined the NSC populations in both niches. We saw a decrease in the quiescent NSC pools in both the SVZ and the SGZ, which corresponded to an increase in intermediate progenitor cells (IPCs) and differentiating neuroblasts. This expansion of proliferative cells closely resembled the NSC phenotypes seen in mice deplete of the NRMT1 target retinoblastoma protein (RB) (Ferguson et al. 2002; Tooley et al. 2010). In RB deficient mice, the expanded populations of IPCs and neuroblasts differentiate into neurons, however, abnormal expression of cell cycle genes prevents these neurons from exiting the cell cycle, inhibits the completion of terminal differentiation, and promotes apoptosis (Ferguson et al. 2002; Andrusiak et al. 2012; Naser et al. 2016).

To understand if the phenotypes seen with loss of NRMT1 could result from misregulation of RB function, we looked at expression of RB target genes in *Nrmt1*^-/-^ mice. During normal neurogenesis, hypophosphorylated RB inhibits E2F-mediated transcription of cell cycle genes and enables both NSC quiescence and terminal differentiation (Andrusiak et al. 2012). Phosphorylation of RB during the proliferative phases of neurogenesis releases this inhibition and promotes RB protein degradation and cell cycle entry (Park et al. 2000). Accordingly, the loss of RB results in an expansion of IPCs and neuroblasts within the niches (Ferguson et al. 2002; Naser et al. 2016). Although these cells migrate and differentiate into mature neurons, they ultimately undergo apoptosis due to a failure to exit the cell cycle and complete terminal differentiation (Andrusiak et al. 2012; Naser et al. 2016). As predicted, we see transcriptional derepression of RB/E2F1 target genes that promote cell cycle entry in *Nrmt1*^-/-^ mice and a corresponding increase in RB phosphorylation and decrease in overall RB protein levels. As observed in RB deficient mice, we also see mature neurons that have migrated but fail to exit cell cycle and ultimately undergo apoptosis. Finally, we confirm previous data in RB-deficient apoptotic neurons that shows apoptosis is not activated by the transcriptional derepression of RB target genes *Puma* and *Apaf1* (Andrusiak et al. 2012) and identify novel derepression of the RB target *Noxa*, an inhibitor of the anti-apoptotic protein MCL-1 (Ploner et al. 2008).

We propose a model where NRMT1 is working through RB to prohibit entry into/promote exit from the cell cycle. Loss of Nα-methylation of RB could result in altered protein/DNA interactions between the N-terminal tail of RB and promoter DNA to produce a conformational change in RB that promotes its phosphorylation, release from E2F1, and subsequent degradation. This would result in the observed loss of RB-mediated transcriptional repression, premature activation of NSC proliferation and differentiation, and failure of mature neurons to exit the cell cycle, ultimately leading to apoptosis. Combined, the depletion of the NSC pool and high rates of neuronal apoptosis would lead to an inability of the niche to replace neurons as they age or become damaged, subsequently leading to the observed neurodegenerative phenotypes. Taken together, these data identify a new role for NRMT1 in NSC development and suggest its regulation of RB contributes to the maintenance of NSC quiescence and their capacity for terminal differentiation.

## Results

### Nrmt1^-/-^ mice exhibit postnatal enlargement of the lateral ventricles and hippocampal neurodegeneration

We have previously seen that *Nrmt1*^-/-^ mice present phenotypes associated with premature aging and phenocopy mouse models deficient in DNA damage repair (Bonsignore et al. 2015b). As an impaired DNA damage response in neurons has been connected to the onset of age-related neurodegenerative disorders, such as Alzheimer’s disease (AD) and amyotrophic lateral sclerosis (ALS) (Madabhushi et al. 2014), we sought to determine whether NRMT1^-/-^ mice exhibit any age-related neural pathologies. We began by staining whole-brain mounted sections of wild type (WT) C57BL/6J mice and C57BL/6J-*Nrmt1*^-/-^ mice with cresyl violet to look for gross morphological defects. In 14-day-old postnatal mice (P14), there were no apparent morphological differences between the WT control and *Nrmt1*^-/-^ brains (Fig. 1A). However, by 6 weeks, *Nrmt1*^-/-^ mice showed a significant enlargement of the lateral ventricles (Fig. 1B,E). This increase became larger at 3 months (Fig. 1C,E) and persisted through 6 months (Fig. 1D,E).

**Figure 1.**
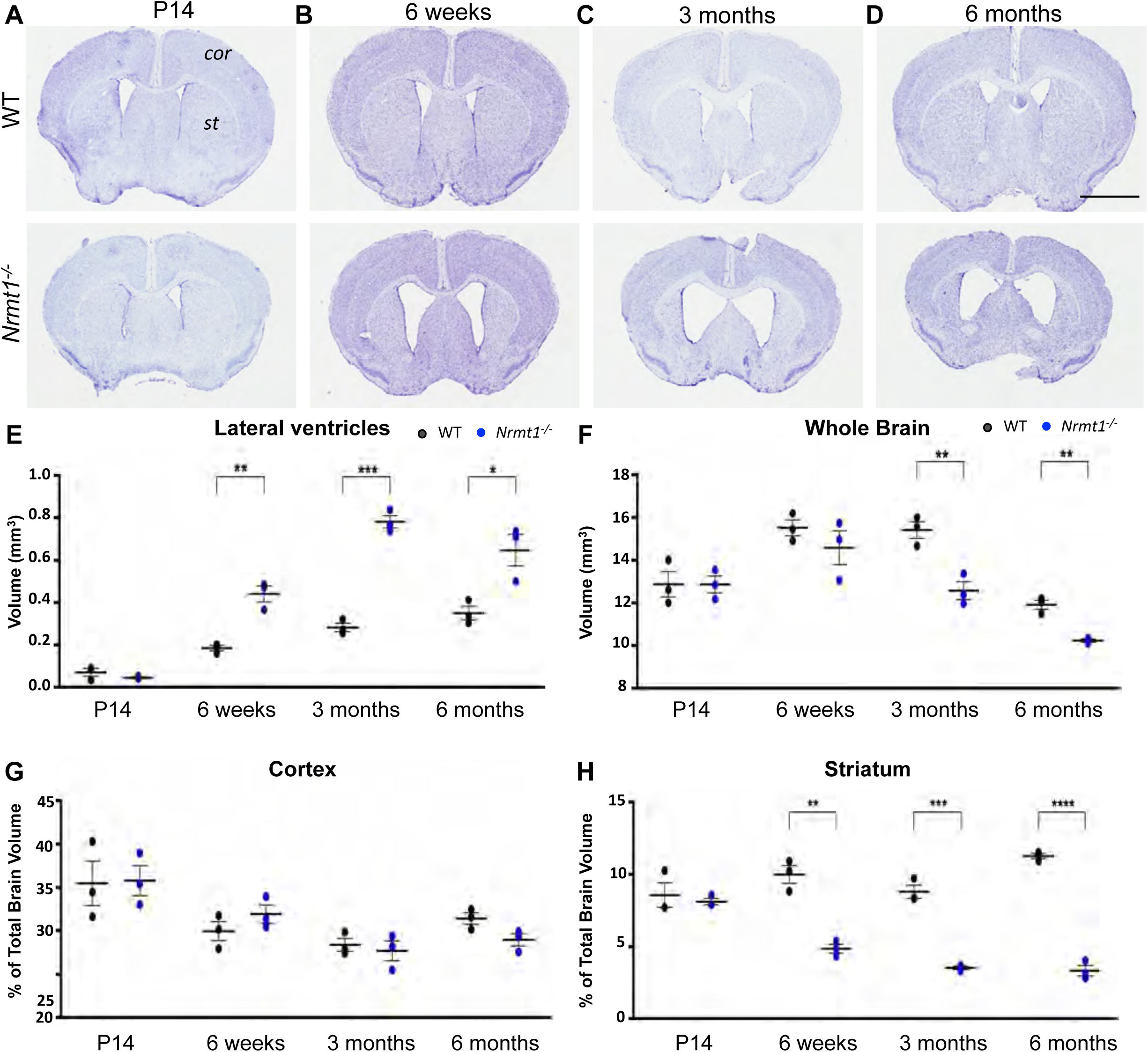
*Nrmt1^-/-^* mice have enlarged lateral ventricles and reduced striatal volume. (*A-D*) Cresyl violet staining showing *Nrmt1^-/-^* mice have enlarged lateral ventricles (black arrow heads) by 6 weeks that persist through 6 months. Quantification of (*E*) lateral ventricle, (*F*) whole brain, (*G*) cortex (*cor*), and (*H*) striatum (*st*) volume. * denotes p<0.05, ** denotes p<0.005, *** denotes p<0.0005, **** denotes p<0.00005 as determined by unpaired t-test, n=3. Error bars represent mean ± SEM. Scale bar = 2 mm.

The lateral ventricles contain cerebrospinal fluid (CSF), which circulates nutrients through the brain and helps remove waste. Ventricle enlargement can be due to global changes in brain volume, where all regions increase proportionally (Horga et al. 2011), or come at the expense of reduced volume in neighboring regions of gray matter (Horga et al. 2011). To determine if enlargement of the lateral ventricles in the *Nrmt1*^-/-^ mice is due to an average increase in brain volume or a loss of neighboring gray matter, volumes of the striatum, cortex, and total brain were calculated. *Nrmt1*^-/-^ mice did not display global increases in brain volume at any age and overall volume actually decreased with age (Fig. 1F). At 6 weeks, when the ventricle volume increased significantly, overall brain volume did not differ and cortical volume also remained similar (Fig. 1F,G). However, the proportional volume of the striatum to total brain volume was already significantly decreased at 6 weeks (Fig. 1H), indicating the lateral ventricle enlargement seen in *Nrmt1^-/-^* mice corresponds to reduced striatal volume. At both 3 and 6 months, *Nrmt1*^-/-^ brains had significantly decreased overall volume (Fig. 1F). The proportional volume of the *Nrmt1*^-/-^ cortex at both these ages showed no difference from WT (Fig. 1G), but the proportional volume of the *Nrmt1*^-/-^ striatum remained significantly different from WT at both 3 and 6 months (Fig. 1H). These data indicate that the lateral ventricle enlargement seen in *Nrmt1*^-/-^ mice occurs concurrently with a decrease in striatal volume.

To next examine the general morphology of the hippocampus, we performed cresyl violet staining of whole-brain mounted sections that contained the hippocampus from P14, 6 weeks, 3 months, and 6 months. The hippocampal circuit is comprised of four main regions, the three *Cornu Ammonis* subfields (CA1, CA2, CA3) and the dentate gyrus (DG). At P14 and 6 weeks, no obvious morphological differences were observed between *Nrmt1*^-/-^ and WT mice (Fig. 2A,B). However, at 3 months, the volumes of both the CA3 and DG are significantly decreased in *Nrmt1*^-/-^ mice, indicating neuronal loss (Fig. 2C,E,F). By 6 months, the volumes of the CA3 and DG are even further decreased, indicating that significant neurodegeneration has occurred by this age (Fig. 2D,E,F). Taken together, these data show that *Nrmt1*^-/-^ mice exhibit both abnormal developmental morphologies and age-related neurodegeneration.

**Figure 2.**
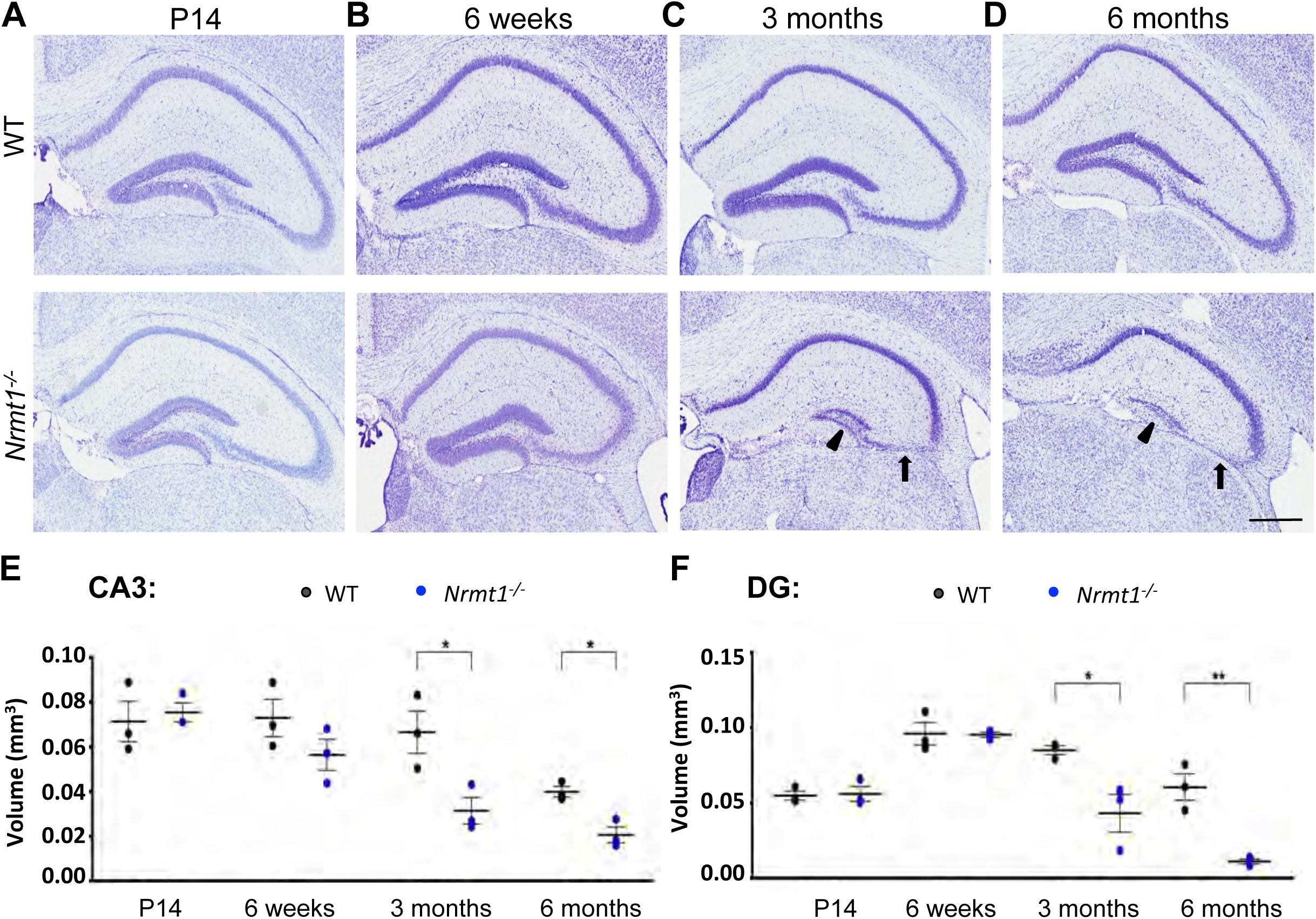
*Nrmt1^-/-^* mice show degeneration of hippocampal neurons. Cresyl violet staining of hippocampal neurons shows similar morphology between *Nrmt1^-/-^* and WT mice at (*A*) P14 and (*B*) 6 weeks. However, at (*C*) 3 months *Nrmt1^-/-^* mice show degeneration of neurons in the CA3 (arrow) and DG (arrowhead). (*D*) By 6 months, neuronal staining is almost completely absent in the CA3 and DG. Quantification of (*E*) CA3 and (*F*) DG volume. * denotes p<0.05 and ** denotes p<0.005 as determined by unpaired t-test, n=3. Error bars represent mean ± SEM. Scale bar = 400 µm.

### Behavioral analysis of Nrmt1^-/-^ mice

Enlargement of the lateral ventricles and/or neurodegeneration of specific brain regions are common pathologies seen in many types of brain disorders including schizophrenia, Down syndrome, autism spectrum disorder (ASD), attention deficit and hyperactivity disorder (ADHD), Alzheimer’s disease (AD), Parkinson’s disease (PD), Huntington’s disease (HD), and Amyotrophic lateral sclerosis (ALS) (Piven et al. 1995; Lyoo et al. 1996; Wright et al. 2000; Guptha et al. 2002; Ishihara et al. 2010; Westeneng et al. 2015). Many of these disorders are associated with memory or motor impairments. Since the hippocampus contributes to many aspects of short-term working memory (Kumaran 2008), long-term declarative memory (Squire and Zola-Morgan 1991), and spatial learning/memory (Burgess 2002), and the striatum is the major input for voluntary motor control (Hikosaka et al. 2000), we wanted to see if *Nrmt1*^-/-^ mice similarly exhibited memory or motor impairments. First, we used a two-trial Barnes maze test to assess short-term spatial and working memory. In this task, mice were given two three-minute training periods to find the correct escape hatch from the maze. Fifteen minutes after the final training trial, the escape hatch was removed and the animals were placed back on the platform. Exploration time at the hole that previously housed the escape hatch (correct hole) was measured and compared to time spent at the other incorrect holes. At 6 weeks and 3 months, *Nrmt1*^-/-^ mice spent significantly less time exploring the correct hole than WT controls during the test trials (Fig. 3A,B). Furthermore, by 6 months, *Nrmt1*^-/-^ mice spent significantly more time at the incorrect holes than the correct hole (Fig. 3C). To summarize the test trial data, a spatial memory index for all ages was calculated by taking the difference between correct versus incorrect hole exploration and dividing the difference by total exploration time. A negative number for the spatial memory index indicates the animal did not remember the escape hatch during the test trial. At all ages, the spatial memory index was significantly decreased in *Nrmt1*^-/-^ mice as compared to the WT controls (Fig. 3D), indicating *Nrmt1*^-/-^ mice have short-term spatial memory deficits.

**Figure 3.**
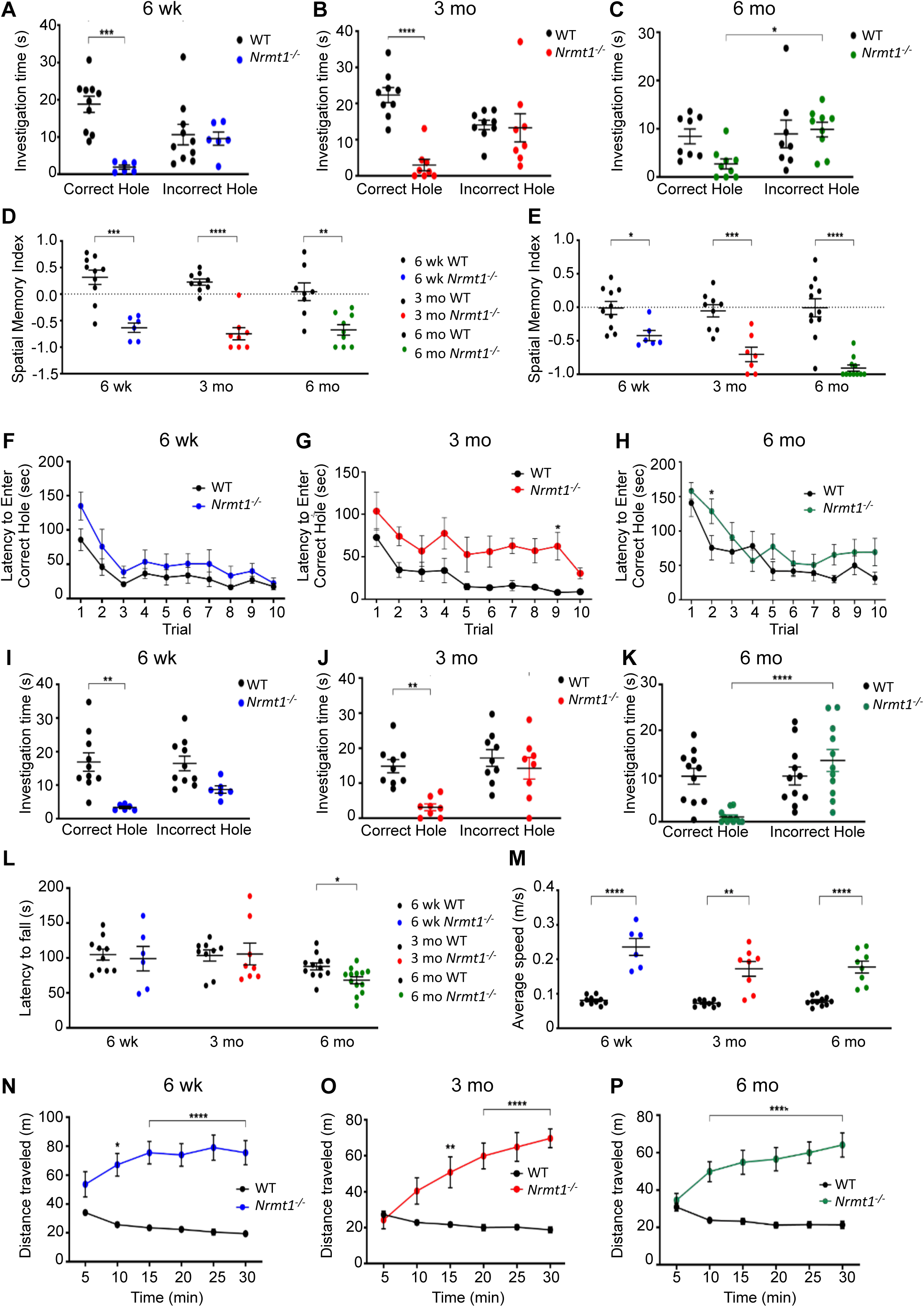
*Nrmt1^-/-^* mice exhibit short and long-term memory impairment and hyperactivity. Two-trial Barnes maze spatial memory tasks show that *Nrmt1^-/-^* mice spend significantly less time exploring the correct hole than WT mice at (*A*) 6 weeks and (*B*) 3 months. (*C*) At 6 months, *Nrmt1^-/-^* mice spend significantly more time at the incorrect holes. (*D*) The spatial memory indexes (SMI) calculated from the two-trial tasks indicate short-term memory loss at all three ages. SMI = (time exploring correct hole – time exploring incorrect holes)/(total exploration time). (*E*) The SMIs calculated from the ten-trial tasks also indicate long-term memory loss at all three ages. Though *Nrmt1^-/-^* mice are able to learn the correct hole when given ten training trials at (*F*) 6 weeks, (*G*) 3 months, and (*H*) 6 months, the ten-trial Barnes maze also shows *Nrmt1^-/-^* mice spend significantly less time exploring the correct hole at (*I*) 6 weeks and (*J*) 3 months and significantly more time at incorrect holes at (*K*) 6 months. (*L*) Rotarod assays show a small motor impairment in *Nrmt1^-/-^* mice at 6 months. Open field assays show *Nrmt1^-/-^* mice travel at (*M*) higher average speeds for (N-P) longer distances than WT mice. * denotes p<0.05, ** denotes p<0.005, *** denotes p<0.0005 as determined by (*A-C, H-J*) two-way anova or (*D, E-G, K, L, M-P*) unpaired t-test, n=6-10. Error bars represent mean ± SEM.

The spatial memory index was significantly decreased in *Nrmt1*^-/-^ mice as compared to the WT controls after a ten-trial Barnes maze test as well (Fig. 3E). Given that two training trials may not have been enough time for the *Nrmt1*^-/-^ mice to learn where the escape hatch was, and to rule out the possibility that these animals have learning impairments rather than memory deficits, a 10-training trial Barnes maze test was also performed. Mice were now given ten three-minute training periods to find the correct escape hatch from the maze. During each trial, the time it took for each mouse to find the correct hole (latency to enter correct hole) was measured. Though the *Nrmt1*^-/-^ mice progressed somewhat slower, by the tenth trial they showed no significant difference from control animals in their ability to locate the escape hole at any age (Fig. 3F-H), indicating they could learn the task. Twenty-four hours after the last training trial, the escape hatch was removed and exploration time at the correct and incorrect holes was measured. Again, at 6 weeks and 3 months, *Nrmt1*^-/-^ mice spent significantly less time exploring the correct hole than WT controls (Fig. 3I,J), and at 6 months, *Nrmt1*^-/-^ mice spent significantly more time at the incorrect holes than the correct hole (Fig. 3K), indicating *Nrmt1*^-/-^ mice also have impaired long-term memory.

The inability of *Nrmt1*^-/-^ mice to locate and explore the correct versus incorrect holes in the Barnes maze could also result from impaired motor activity. To test motor performance in *Nrmt1*^-/-^ mice, rotarod tests were performed. Mice were placed on an accelerating rotarod for three training trials, followed by two test trials. The amount of time each animal stayed on the rotarod before falling (latency to fall) was measured. There was no significant difference in the latency to fall between *Nrmt1*^-/-^ mice and WT controls at 6 weeks or 3 months (Fig. 3L). At 6 months, there was a small, yet significant, decrease in motor performance in the *Nrmt1*^-/-^ mice (Fig. 3L). However, this small difference at 6 months on the rotarod cannot account for the impaired performance of *Nrmt1*^-/-^ mice at earlier ages in the Barnes maze. To confirm these results, we also performed open field assays to test the motor activity of *Nrmt1*^-/-^ mice. Surprisingly, at all ages, *Nrmt1*^-/-^ mice moved at a significantly higher average speed and traveled significantly longer distances than WT controls (Fig. 3M-P), indicating that *Nrmt1*^-/-^ mice do not have any significant motor impairments, although they do exhibit locomotor hyperactivity.

This hyperactive phenotype may be partly explained by striatal function. The striatum is a crucial input for motor control, with dopamine D1 receptors involved in voluntary movement and dopamine D2 receptors responsible for inhibition of movement (DeLong 1990; Graybiel 1990). An increase in D1 activity or decrease in D2 function has been shown to facilitate hyperactive phenotypes in mice (Sano et al. 2003; Li and Zhou 2013). Given the decreased striatal volume in our mouse, selective death of D2 expressing neurons may explain the observed hyperactivity. Neuropathological studies of HD show a similar phenomenon, with almost exclusive loss of inhibitory striatal neurons during the early stages of chorea (Chen et al. 2013). Striatal inhibitory neurons are more excitable, and it is hypothesized this makes them more susceptible to abnormal neurotransmitter levels or receptor dysfunction (Chen et al. 2013). Our data demonstrate that the inability of *Nrmt1*^-/-^ mice to explore the correct hole in the Barnes maze is due to memory and not motor impairments. The observed memory impairments begin before any obvious hippocampal degeneration, indicating that similar to HD (Chen et al. 2013), neuronal dysfunction is preceding cell death.

### Molecular analysis of the neural stem cell niches

The morphological changes seen in the lateral ventricles and hippocampus and the corresponding behavioral phenotypes suggested a potential misregulation of adult NSC neurogenesis, as the two postnatal NSC niches are found in the subventricular zone (SVZ) of the ventricles and the subgranular zone (SGZ) of the DG (Obernier and Alvarez-Buylla 2019). The SVZ contains the radial glia-like stem cells (RGLs) that produce both ependymal cells (E1 cells) and neural stem cells (named B1 cells in the SVZ) (Redmond et al. 2019). These RGLs account for the majority of proliferating cells in the ventricular zone (Bátiz et al. 2011). E1 cells are first produced between embryonic day 14 (E14) and embryonic day 16 (E16), mature during the first postnatal week, and become postmitotic after maturation (Spassky et al. 2005; Bátiz et al. 2011). Postnatal production of new E1 cells results from proliferation of RGLs and their initiation of ependymogenesis (Bátiz et al. 2011). Increased proliferation and ependymogenesis of RGLs has been shown to sustain ventricle enlargement in mouse model of hydrocephalus (Bátiz et al. 2011). To determine if increased amounts of cell proliferation were preceding ventricle volume enlargement in *Nrmt1*^-/-^ mice, brain sections were immunostained for the proliferation marker Ki-67. Remarkably, there was a significant increase in cell proliferation around the SVZ at P14 (Fig. 4A,C). These data show that increased cell proliferation in the ventricular walls precedes ventricular enlargement and suggest abnormal activation of RGL proliferation in *Nrmt1*^-/-^ mice.

**Figure 4.**
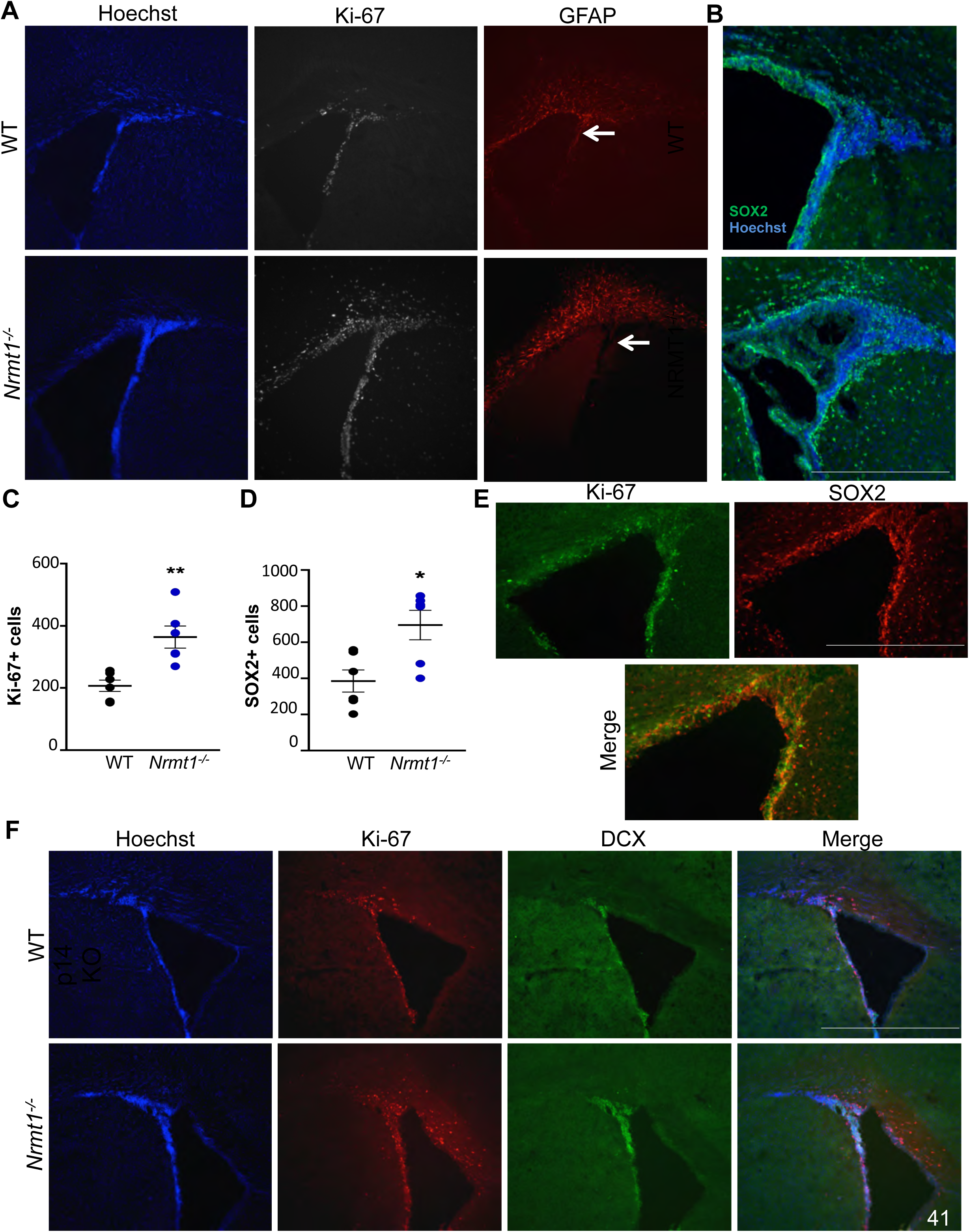
At P14, *Nrmt1^-/-^* mice exhibit a depletion of the RGL pool in the SVZ and a corresponding expansion of the IPC and neuroblasts pools. (*A,C*) Though there is a significant increase of Ki-67 positive cells in the SVZ of *Nrmt1^-/-^* mice, GFAP staining is absent near the SVZ (arrows) and does not overlap with regions of increased Ki-67 staining. There is a also a significant increase in (*B,D*) SOX2 positive IPCs and (*F*) increased Doublecortin (DCX) immunostaining. (*E,F*) Ki-67 staining patterns overlap with both SOX2 and DCX. * denotes p<0.05 and ** denotes p<0.005 as determined by unpaired t-test, n=6. Error bars represent mean ± SEM. Scale bar = 1000 µm.

To determine if these proliferating cells in *Nrmt1*^-/-^ mice were RGLs, we stained ventricular sections of P14 mice with antibodies against GFAP (a marker of both RGLs and astrocytes). GFAP staining near the SVZ was absent in *Nrmt1*^-/-^ mice and did not overlap with the regions of increased Ki-67 staining, indicating these proliferating cells are not RGLs (Fig. 4A). As the proliferating cells could be descendants of RGLs that have begun differentiation, we also stained ventricle sections of P14 mice with antibodies against SOX2, a marker of cells in the B1 lineage, which also arise from RGLs. We found a significant increase in SOX2 positive (SOX2+) cells in and around the SVZ in *Nrmt1*^-/-^ mice as compared to WT (Fig. 4B,D). Since GFAP is a marker for RGLs and SOX2 marks both RGLs and intermediate progenitor cells (IPCs) (Hutton and Pevny 2011), the decrease in GFAP staining and increase in SOX2+ cells in *Nrmt1*^-/-^ mice indicate a shift from normal basal neurogenic activity to an aberrant expansion of IPCs. While the SOX2 and Ki-67 patterns were very similar, there was not complete overlap (Fig. 4E). To determine if the differentiation of these NSCs is proceeding past the IPC stage to form neuroblasts and if these neuroblasts are also actively proliferating, co-staining with Ki-67 and the neuroblast marker Doublecortin (DCX) was performed. There was a detectable increase in DCX staining in the ventricle of *Nrmt1*^-/-^ mice, which corresponded to regions of increased Ki-67 staining (Fig. 4F), indicating abnormal neuroblast proliferation is also occurring in the SVZ.

We next looked to see if similar NSC misregulation was occurring in the SGZ. The DG is composed of three main layers; the outer molecular layer; the granule cell layer (GCL), which contains the subgranular zone (SGZ); and the hilus/polymorph layer, which is located between the GCLs. The DG normally undergoes lifelong neurogenesis (Su et al. 2019), with NSCs from the SGZ ultimately differentiating into the granule neurons that populate the GCL (Goncalves et al. 2016). To look at NSC proliferation in the DG, we again immunostained for Ki-67. Similar to what was seen in the SVZ, at P14 there were significantly more Ki-67 positive (Ki-67+) cells in the SGZ of *Nrmt1*^-/-^ mice than in WT controls, and only *Nrmt1*^-/-^ mice had populations of proliferating cells in the hilus (Fig. 5A,C). To determine if the increase in proliferation found within the SGZ and hilus was due to an expansion of RGLs or cells proceeding down the NSC lineage, P14 hippocampal sections were stained for GFAP, SOX2, and DCX. While WT mice had organized GFAP+ cells lining both sides of the SGZ, *Nrmt1*^-/-^ mice lacked GFAP staining in the SGZ (Fig. 5A), indicating a depletion of RGLs. However, there was a significant increase in SOX2+ cells around the SGZ of *Nrmt1*^-/-^ mice, and only *Nrmt1*^-/-^ mice had populations of SOX2+ cells in all three layers of the DG (Fig. 5B,D). Similar to the SVZ, not all Ki-67+ cells co-stained with SOX2 (Fig. 5E), but there was increased DCX staining in the SGZ that also overlapped with the Ki-67 pattern (Fig. 5F). Taken together, these data indicate abnormal proliferation of IPCs and neuroblasts in both stem cell niches of *Nrmt1*^-/-^ mice.

**Figure 5.**
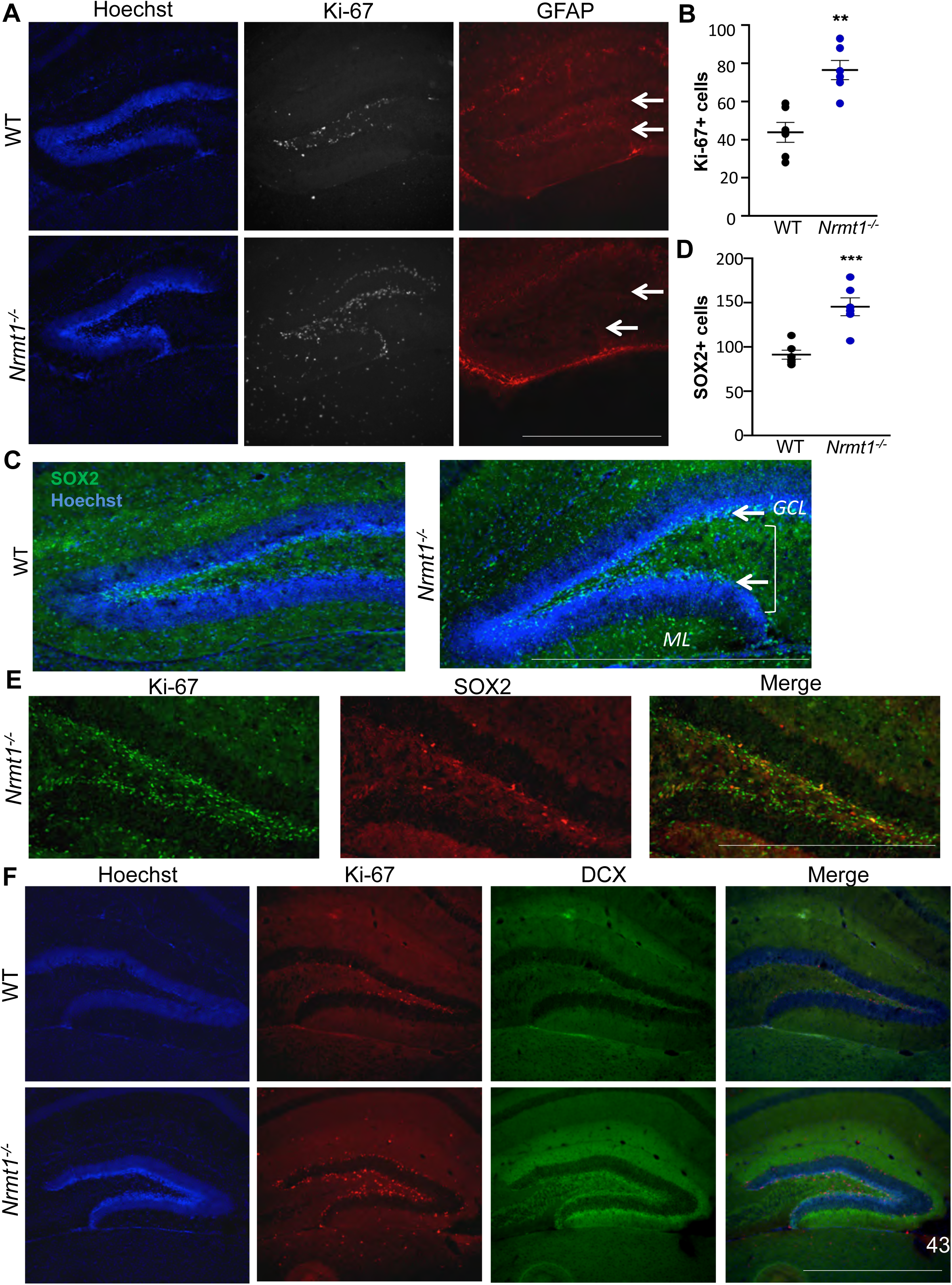
At P14, *Nrmt1^-/-^* mice also exhibit a depletion of the RGL pool and an expansion of IPCs and neuroblasts in the SGZ. (*A,B*) As seen in the SVZ, there is a significant increase of Ki-67 positive cells in the SGZ of *Nrmt1^-/-^* mice. Immunostaining also shows GFAP staining specifically absent from the SGZ (arrows) and no overlap between GFAP and Ki-67 staining patterns. (*C,D*) SOX2 positive cells are also significantly increased in the SGZ (arrows), granule cell layer (GCL), molecular layer (ML) and hilus (bracket) of *Nrmt1^-/-^* mice, and (*E*) there is overlap between the SOX2 and Ki-67 staining patterns. (*F*) Regions of increased DCX immunostaining in the dentate gyrus of *Nrmt1^-/-^* mice also correspond to regions of increased Ki-67 immunostaining. ** denotes p<0.005 and *** denotes p<0.0005 as determined by unpaired t-test, n=6. Error bars represent mean ± SEM. Scale bar = 1000 µm.

### Functional analysis of retinoblastoma protein in Nrmt1^-/-^ mice

The expansion of IPC and neuroblast populations seen in *Nrmt1*^-/-^ mice was extremely reminiscent of phenotypes seen in mice deficient for the NRMT1 target protein RB. Mice with a conditional knockout of RB in the telencephalon exhibit abnormal proliferation of cells that co-stain with TuJ1, a marker of neuroblasts/early neurons, indicating that even though the NSC cell cycle is misregulated, they can still commit to a neuronal fate and initiate differentiation (Ferguson et al. 2002). Temporal loss of RB in mouse NSCs also results in increased proliferation of IPCs and neuroblasts and increased neuroblast migration through the rostral migratory stream (RMS) (Naser et al. 2016). Once the migratory neuroblasts reach the olfactory bulb, they are able to differentiate into mature interneurons, but these interneurons are gradually lost over time (Naser et al. 2016). In culture, acute removal of RB results in neurons that stain for the proliferative marker Ki-67, abnormally express cyclin genes, and begin to undergo apoptosis (Andrusiak et al. 2012), indicating that the delayed loss of RB deficient neurons *in vivo* results from their inability to exit the cell cycle and complete terminal differentiation (Andrusiak et al. 2012).

RB normally acts to inhibit the cell cycle through its repression of E2F-mediated transcription of cell cycle regulatory genes, including many cyclins, cell division cycle genes, and even *E2f1* itself (Neuman et al. 1995; DeGregori 2002; White et al. 2005). To monitor if RB function is disrupted in *Nrmt1*^-/-^ mice, we looked at mRNA expression of *Cyclin A2*, *Cyclin E2*, and *E2f1*. Consistent with expression patterns found with RB deficiency (Andrusiak et al. 2012; Ghanem et al. 2012), we found that expression of all three cell cycle genes is increased in *Nrmt1*^-/-^ mice (Fig. 6A). We then used western blots to see if this transcriptional derepression of RB target genes corresponded to changes in RB phosphorylation or protein levels. In *Nrmt1*^-/-^ mice, phosphorylation of RB at Ser795 significantly increases, while overall RB protein levels significantly decrease (Fig. 6B,C), indicating that loss of Nα-methylation triggers the release of RB from E2F proteins (Park et al. 2000; DeGregori 2002).

**Figure 6.**
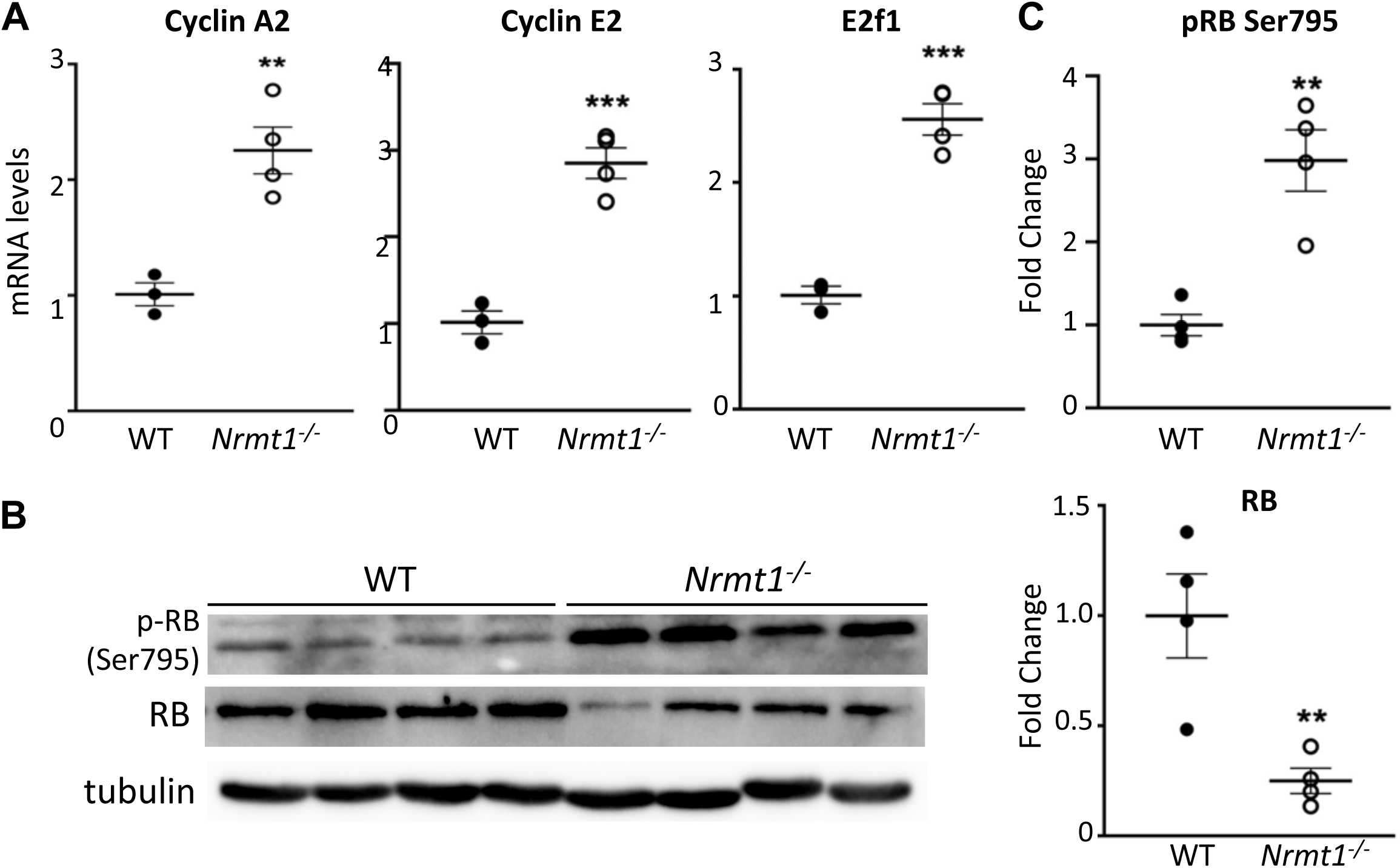
*Nrmt1^-/-^* mice show misregulation of RB function. (*A*) mRNA levels of RB target genes *Cyclin A2*, *Cyclin E2*, and *E2f1* are significantly increased in *Nrmt1^-/-^* mice. (*B*) In addition, RB phosphorylation at serine 795 (p-RB Ser795) significantly increases but overall RB protein levels significantly decrease. (*C*) Quantification of western blots. All samples normalized to tubulin loading control. ** denotes p<0.005 and *** denotes p<0.0005 as determined by unpaired t-test, n=3-4. Error bars represent mean ± SEM.

To determine if the activation of cell cycle genes resulted in an inability of *Nrmt1*^-/-^ neurons to exit the cell cycle, we co-stained ventricle/striatal sections from P14 mice with NeuN (a marker of mature neurons) and Ki-67. Ki-67 staining was restricted to the SVZ in WT mice and no mature neurons in the striatum co-stained with NeuN and Ki-67 (Fig. 7A). However, in *Nrmt1*^-/-^ mice, Ki-67+ cells were present in both the SVZ and neighboring striatum and many of the Ki-67+ cells in the striatum co-stained with NeuN (Fig. 7A). These data indicate loss of NRMT1, similar to loss of RB, is preventing neuronal exit from the cell cycle. The population of NeuN/Ki-67 double positive cells in *Nrmt1*^-/-^ mice persisted through 3 weeks, decreased by 4 weeks, and was primarily gone by 6 weeks (Fig. 7B). Double positive cells were not seen in WT mice at any age (Supplemental Figure 1A). To see if and when the double positive cells in *Nrmt1*^-/-^ mice undergo apoptosis, we stained ventricle/striatal sections for the apoptotic marker cleaved caspase-3. Very little cleaved caspase-3 staining is visible at P14 or 3 weeks in WT or *Nrmt1*^-/-^ mice (Fig. 7B, Supplemental Figure 1A-B). However, while WT mice remain devoid of cleaved caspase-3 staining at 4 and 6 weeks (Supplemental Figure 1A), punctate cleaved caspase-3 staining begins to appear in the striatum of *Nrmt1*^-/-^ mice by 4 weeks and is prominent by 6 weeks (Fig. 7B). These data indicate that, as with RB deficiency, neurons in *Nrmt1*^-/-^ mice are remaining in the cell cycle, failing to complete terminal differentiation, and ultimately undergoing a delayed apoptosis.

**Figure 7.**
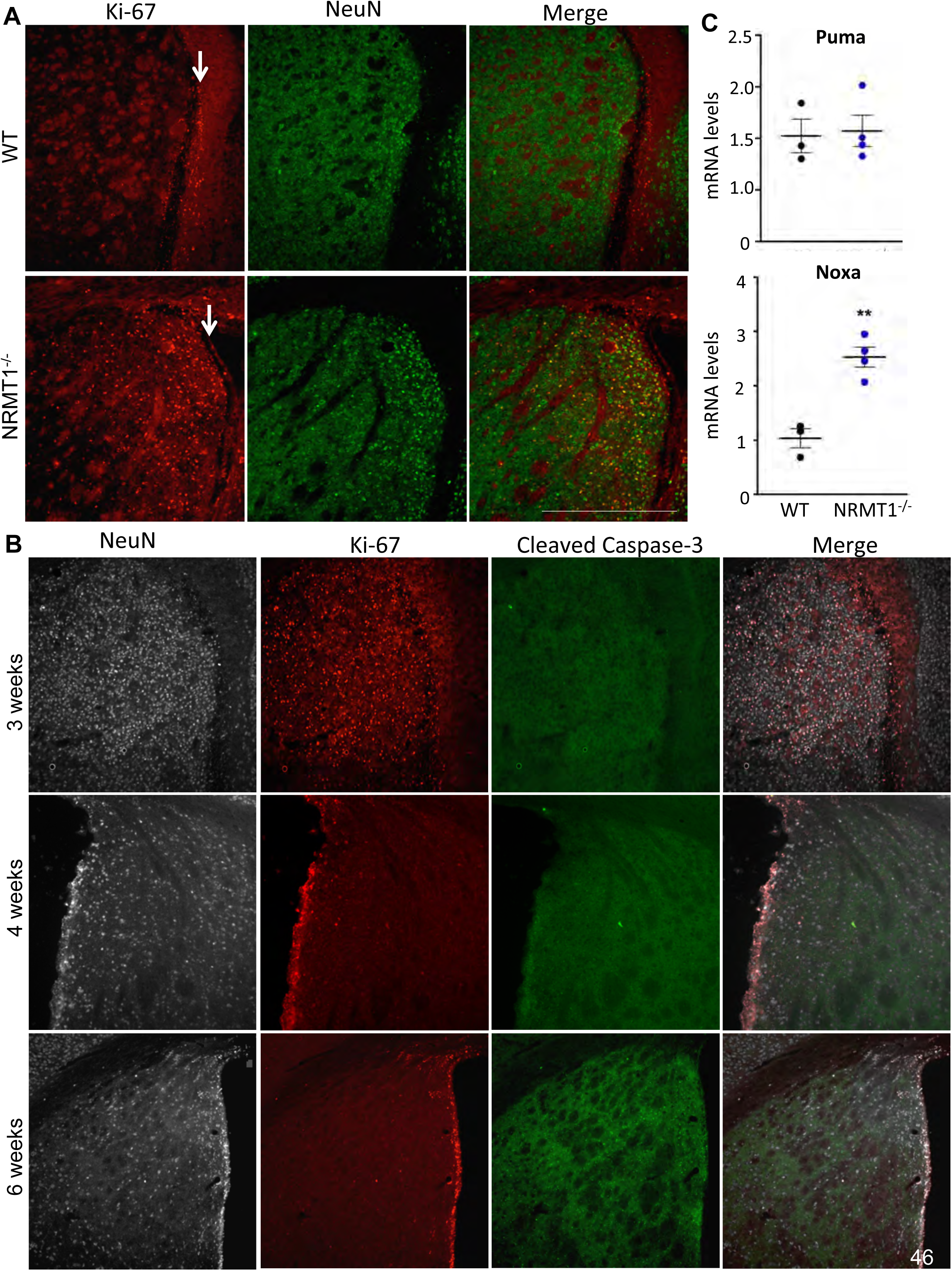
Neurons in *Nrmt1^-/-^* mice fail to exit the cell cycle and ultimately undergo NOXA-mediated apoptosis. (*A*) At P14, Ki-67 staining (red) is restricted to the SVZ (arrow) in WT mice. In *Nrmt1^-/-^* mice, Ki-67+ cells are found in the SVZ and neighboring striatum and many co-stain (yellow) with NeuN (green), a marker of mature neurons. (*B*) Ki-67 (red) and NeuN (gray) double positive neurons (pink) persist in *Nrmt1^-/-^* mice through 3 weeks of age but begin to decrease by 4 weeks and are mostly gone by 6 weeks. Punctate cleaved caspase-3 staining (green) begins to appear by 4 weeks and is prominent by 6 weeks. (*C*) qRT-PCR analysis indicates apoptosis is not driven through increased expression of the RB target *Puma* but instead by the RB target *Noxa*. ** denotes p<0.005 as determined by unpaired t-test, n=3-4. Error bars represent mean ± SEM. Scale bar = 1000 µm.

It has been hypothesized that the neuronal apoptosis seen with RB loss is triggered by transcriptional derepression of the RB/E2F target genes *Puma* and *Apaf1*, but increased expression of these genes was not detected in RB-deficient neurons grown in culture (Andrusiak et al. 2012). We assayed *Puma* and *Apaf1* transcript levels in our model. *Apaf1* transcripts were not detectable in either WT or *Nrmt1*^-/-^ ventricle/striatal samples (data not shown), and there was no detectable difference in *Puma* levels (Fig. 7C). However, RB is also a transcriptional regulator of the apoptotic driver *Noxa* (Hershko and Ginsberg 2004). While PUMA is a direct activator of BAX-induced apoptosis, NOXA is an inhibitor of anti-apoptotic proteins (Hollville et al. 2019), and they have been shown to differentially participate in dual pathways of p53- mediated apoptosis induction (Shibue et al. 2006). Interestingly, we do see a significant increase in *Noxa* expression in *Nrmt1*^-/-^ mice (Fig. 7C), indicating PUMA and NOXA may also play differential roles in RB-mediated apoptosis induction, with NOXA signaling predominating in cells that fail to undergo postmitotic cell cycle arrest. Taken together, our data suggest that misregulation of RB-dependent cell cycle arrest in NSCs contributes to the morphological and behavior phenotypes seen in *Nrmt1*^-/-^ mice.

## Discussion

Neural stem cells are somatic stem cells that have the ability to both self-renew and undergo neurogenesis. While NSCs were originally thought to be primarily embryonic and play a pivotal role in CNS development (Alvarez-Buylla and Lim 2004), it has now been widely recognized that NSCs are also present and functional in the postnatal brain (Obernier and Alvarez-Buylla 2019). Postnatal neurogenesis promotes plasticity and likely contributes to olfactory learning and memory formation, as well as provides a source for replacement of cells lost to turnover or injury (Kitabatake et al. 2007; Ma et al. 2009; Nissant et al. 2009). Here we show for the first time that loss of the N-terminal methyltransferase NRMT1 results in the misregulation of postnatal NSCs. Early depletion of quiescent NSCs and expansion of both the IPC and neuroblasts pools is followed by morphological and behavioral abnormalities, and ultimately, neurodegeneration. We propose the observed striatal and hippocampal neurodegeneration results from depletion of the NSC pool and an increase in apoptotic neurons that fail to exit the cell cycle and undergo RB-mediated terminal differentiation.

NRMT1 (also known as METTL11A) is a member of the Methyltransferase like (METTL) family of methyltransferases, and while we show the first identified role for NRMT1 in NSC fate determination, there is an abundance of recent data showing roles for different members of the METTL family in stem cell development. The *N*^6^-methyladenosine (m^6^A) methyltransferases METTL3 and METTL14 regulate the self-renewal of adult hematopoietic stem cells by promoting the expression of genes that maintain quiescence (Yao et al. 2018). Additionally, METTL14 has been shown to regulate embryonic NSCs proliferation and differentiation (Wang et al. 2018). METTL3 has been shown to control adipogenic differentiation through regulation of the JAK/STAT pathway (Yao et al. 2019) and regulate heterochromatin formation in embryonic stem cells (Xu et al. 2021). The more recently discovered m^6^A methyltransferase METTL5 is needed for embryonic stem cell pluripotency (Ignatova et al. 2020). Its loss results in decreased global translation rates and lowered differentiation potential (Ignatova et al. 2020). METTL17, a mitochondrial RNA methyltransferase, regulates the stability of 12S mt-rRNA and its associated proteins, and is required for differentiation of mouse embryonic stem cells (Shi et al. 2019). The closest sequence homolog to NRMT1, NRMT2 (also known as METTL11B), is an Nα-monomethylase, and its transcript levels have been shown to increase during osteocytic and myogenic differentiation (Hong et al. 2020). We have seen a similar upregulation of NRMT1 transcript levels during myogenic differentiation (unpublished data).

Similar to other METTL family members, our data indicate that NRMT1 normally acts to constrain entry into or promote exit from the cell cycle. This may be why embryonic development in *Nrmt1*^-/-^ mice looks morphologically unaffected. In contrast to adult NSCs, which spend a significant amount of time in G0, embryonic NSCs have high proliferation rates (Urbán and Guillemot 2014). If NRMT1 hinders entry into the cell cycle, its levels would be expected to be at low levels during embryogenesis and increase as embryonic NSC proliferation culminates and differentiation begins. RNA-seq data from mouse neural tissue support this model, as they show low NRMT1 mRNA levels during embryogenesis that drastically increase at P7 (Bennett et al. 2016; Clarke et al. 2018). P7 is an important developmental time point in mice, as the early developmental stages have been completed and areas such as the DG are formed but just beginning to mature into neuronal layers (Gilley et al. 2011). These data suggest that NRMT1 expression increases as NSCs begin to leave their proliferative phase, concentrate in their neurogenic stem cells niches, and take on regenerative instead of developmental roles. Future single-cell RNA-sequencing (scRNA-seq) analysis will be useful for monitoring NRMT1 expression during the stages of NSC differentiation (Mizrak et al. 2019) and determine if its levels normally decrease during IPC and neuroblast proliferation, as our model also predicts.

Here we show that the neural phenotypes seen in *Nrmt1*^-/-^ mice closely resemble those seen in mice deficient for the NRMT1 target RB (Tooley et al. 2010). RB is a well-known tumor suppressor protein that inhibits cell cycle progression and apoptosis (Dick and Rubin 2013) but has also been show to regulate neuroblast differentiation and migration (Ghanem et al. 2012). As Nα-methylation regulates DNA/protein interactions, we propose a model where loss of NRMT1 disrupts the interaction of the N-terminal RB tail with DNA at target promoters and alters the conformation of the RB N-terminal domain (RbN). When RbN interacts with the phosphorylated RB interdomain linker (RbIDL), they are able to bind the RB pocket domain and displace E2F (Burke et al. 2010). We propose loss of Nα-methylation of the RB tail promotes interaction between RbN and RbIDL and displacement of RB from E2F. This leads to increased phosphorylation and degradation of RB and subsequent activation of E2F target genes. At early stages, activation of cell cycle genes results in abnormal activation and proliferation of the quiescent NSC population. In later stages, it prohibits neurons from exiting the cell cycle and triggers apoptosis. Our model suggests that it is the combination of a depleted NSC pool and increased neuronal apoptosis that produces the neurodegenerative phenotypes seen in *Nrmt1*^-/-^ mice.

Though our data indicate RB misregulation is a strong phenotypic driver in *Nrmt1*^-/-^ mice, NRMT1 has a variety of other verified or predicted targets known to play a role in NSC development. The Wnt signaling pathway plays dual roles during NSC development, and the β- catenin inhibitor ICAT, is a predicted target of NRMT1. ICAT inhibits transcription of β-catenin target genes by disrupting its interaction with the Tcf/Lef-family of transcription factors (Daniels and Weis 2002). During embryonic development, Wnt signaling first fosters proliferation of NSCs, while later it promotes differentiation of NSCs and migration to the SVZ (Marinaro et al. 2012). The role of Wnt signaling can be modulated in this way by interactors of β-catenin that are differentially expressed or modified during embryogenesis and later development (Marinaro et al. 2012). One such interactor, HIPK1, is low during embryogenesis to allow β-catenin to promote proliferation and increases in NSCs in the SVZ to help β-catenin promote exit from the cell cycle and differentiation (Marinaro et al. 2012). NRMT1 could function similarly to HIPK1 through its regulation of ICAT. The transcription factor SOX5, is also a predicted target of NRMT1. SOX5 plays a role in regulating neuroblast migration and post-migratory differentiation, and its loss results in immature neuronal differentiation (Kwan et al. 2008). The verified NRMT1 target SET has also been show to play a role in regulating the rearrangement of chromatin architecture during the transition between pluripotency and differentiation (Bui et al. 2019). While we see RB misregulation in *Nrmt1*^-/-^ mice, and both RB and NRMT1 deficient mice share many similar phenotypes, it is likely other targets are contributing to *Nrmt1*^-/-^ neural phenotypes and future scRNA-seq will help elucidate these targets as well.

We have shown the first role for NRMT1 in neural development, neurodegeneration, and NSC regulation. Loss of NRMT1 results in abnormal lateral ventricle enlargement, striatal and hippocampal neurodegeneration, cognitive and motor impairments, and misregulation of NSC development. However, these phenotypes result from constitutive NRMT1 loss, and it will be interesting to see if acute NRMT1 loss and subsequent restoration respectively stimulate NSC proliferation and allow exit of differentiating neurons from the cell cycle. During both neurodegenerative diseases and stroke, the burden of neuronal loss is high and an exhausted or limited pool of stem cells is unable to effectively replace the lost neurons (Holvoet et al. 2016). Understanding if short-term NRMT1 inhibition could stimulate NSC proliferation and subsequent replacement of damaged neurons in these disease states could lead to a novel way of harnessing the potential of endogenous NSCs as therapeutic agents (Sugaya and Vaidya 2018; Huang and Zhang 2019).

## Materials and methods

### Maintenance of Nrmt1^-/-^ mice

To produce the homozygous C57BL/6J- *Nrmt1*^-/-^ mice used in the described experiments, heterozygous C57BL/6J- *Nrmt1*^+/-^ females are mated to homozygous C57BL/6J- *Nrmt1*^-/-^ males because homozygous null females are sterile. Homozygous progeny are detected through genotyping. All progeny over 21 days of age are ear punched for identification and genotyped as previously described (Bonsignore et al. 2015b). Progeny less than 21 days are genotyped through tail clips after euthanasia. Both sexes of mice are used in these studies. Mice not needed for experiments are euthanized by inhalation of carbon dioxide in a clear container followed by cervical dislocation. Mice needed for histological brain sections are euthanized by perfusion with 4% PFA following anesthesia. Both the breeding and euthanasia protocols are approved by the State University of New York at Buffalo Animal Care and Use Committee.

### Barnes Maze

To assess spatial learning and memory, the Barnes maze test was performed with minor modifications (Pompl et al. 1999). Briefly, animals were habituated for 5 minutes (mins) on a circular platform that contained 8 equally spaced holes. After habituation, the animals were placed back on the platform for either 2 or 10 training (learning) acquisition trials. During the training phase, an escape hole was present for the animal to enter. A bright light above the platform was used as a weak aversive stimulus to encourage the animal to escape during each training trial. Each animal had up to 3 mins to navigate the platform and find the escape hole. Unique shapes and colors on 4 sides of the maze were used as spatial cues to help the animal locate the escape hole. If the animal was unable to locate the correct escape hole in the 3 min training trial, the animal was gently guided to the correct hole. The learning interval between each animal was 5 min. After 2 or 10 training trials, a test trial was administered 15 min or 24 hours after the last training trial, respectively. During the test trial, animals were placed back on the platform for 3 min where correct (T1) and incorrect (T2) hole exploration was measured. Spatial memory index was calculated by: (T1-T2) / (T1+T2).

### Open field assay

To test locomotor activity, animals were placed in a transparent plastic cage (40 x 40 x 30 cm) where they were allowed to freely move around for 30 min. ANYMAZE tracking software (Stoelting, Wood Dale, IL) was used to measure total distance traveled and average speed. Quantification of 5 min bins was used in analysis.

### Rotarod assay

To measure motor function, animals were placed on an accelerating rotarod (SD Instruments, San Diego, CA) as previously described with minor adaptations (Dunham and Miya 1957). The rotarod machine was programmed to accelerate from 4 revolutions per minute (RPM) to 40 RPMs over a 5 min trial. In order for the animal to stay on the accelerating rotating cylinder, they must increase their motor output to keep up with the cylinder. To prevent injury after falling, thick foam was used to catch the animals. Three training trials were used to familiarize the animal to the test. If they fell off the apparatus within the first 30 sec of the practice sessions, they were placed back on the cylinder. 10 min after the last practice session, two test trials were administered where the latency to fall off the rotarod machine was measured. The average latency of fall between the two test trials was used for analysis.

### Cresyl violet staining

Tissue sections were rehydrated in 100% ethanol (EtOH) and 95% EtOH for 2 min before staining. Sections were incubated in 0.01 % cresyl violet solution containing glacial acetic acid for 5 min. Tissue was then dehydrated in a series of EtOH steps (70% EtOH, 95% EtOH, and 100% EtOH) for 2 min before clearing in two passes of xylene for 5 min. Sections were mounted in Permount mounting media (Thermo Fisher Scientific) and coverslips were put on and sealed. Slides were allowed to dry overnight. Slides were scanned and digitized using the Aperio ScanScope whole slide scanner (Leica Biosystems) before further analysis. Three animals were used per each cresyl violet quantification, as the poor overall health of *Nrmt1*^-/-^ mice limited sample availability.

### QuPath Volume Analysis

To assess brain volume of specific regions, two sets of slides were created to represent the whole brain (8 sections per set, each spaced 300 µm apart). As a result, the first set of slides contained the rostral half and the second set covered the caudal half of the brain. One full set of cresyl violet stained slides for each animal was scanned using the Aperio ScanScope whole slide scanner (Leica Biosystems) with the 20x objective. Each digitized file was uploaded onto QuPath (version 0.1.2) where they were further processed. Each section containing the brain region of interest was manually traced using the polygon tool. The mouse brain coronal reference of the Allen Brain Atlas was used as a guide for a more accurate trace of the different brain regions. After each trace was completed, an area (µm^2^) given by QuPath was used to calculate the volume of each tissue section. To calculate the volume of a traced region, the formula: Area of trace (µm^2^) x thickness of section (30 µm) = volume of traced area (µm^3^) was determined for each section and brain region. To calculate total volume of a given region, the sum of all volumes calculated for that region were added to give the final volume.

### Immunohistochemistry

Mice were anesthetized and transcardially perfused with 1% PBS followed by 4% paraformaldehyde (PFA) before brain removal. Brains were post-fixed in 4% PFA for 72 hours and then cryoprotected in a 30% sucrose solution for 1 week. 30 µm coronal slices were cut on a Microm HM 525 cryostat (FisherScientific) and attached to Superfrost plus slides (FisherScientific). Slides were stored in a -80° C freezer before being processed. Briefly, sections were washed with PBS and blocked in 1x PBS containing 2% Triton X-100 and 5% horse serum for 1 hour at room temperature (RT). Sections were incubated overnight at 4° C with the following primary antibodies in 1X PBS containing 2% Triton X-100 and 2% horse serum: mouse anti-NeuN (EMD Millipore), rat anti-Ki-67 (Invitrogen), rabbit anti-Ki67 (Cell Signaling Technologies), rabbit anti-Sox2 (EMD Millipore), rabbit anti-cleaved caspase-3 (Cell Signaling Technologies), rabbit anti-Doublecortin (Abcam), rabbit anti-RB (Cell Signaling Technologies), rabbit anti-phospho RB (Ser 795) (Cell Signaling Technologies), or mouse anti-GFAP (Invitrogen). The following day, sections were washed 3 times with 1x PBS and incubated for 1 hour at RT in the following secondary antibodies in 1x PBS containing 2% Triton X-100 and 2% horse serum: Goat anti-rabbit Alexa Fluor 488 or 594 (Invitrogen), Goat anti-mouse Alexa Fluor 594 or 647 (Invitrogen), or Donkey anti-rat Alexa Fluor 488 or 594 (Invitrogen). Sections were counterstained with Hoechst 33342 (AnaSpec Inc.), washed 3 times in 1x PBS followed by a quick rinse in distilled water, and dried at 60° C for 5 mins. Slides were mounted in vectashield fluorescent anti-fading mounting media (Vector Laboratories). Coverslips were added and sealed and slides were visualized using a Cytation 5 Multi-Mode reader, a Leica DMi8 fluorescence microscope, or a Leica TCS SP8 spectral confocal. All images captured were at identical regions and exposures between genotypes.

### Quantitative Real-time PCR Analysis

Lateral ventricles containing the adjacent striatum was collected from 6 week WT and *Nrmt1*^-/-^ mice and total RNA was lysed and extracted using TRIzol (Life Technologies) and chloroform. 1µg of RNA was used as a template to create cDNA through the SuperScript III first-strand synthesis system (Invitrogen). 2µg of cDNA was then used with SYBR Green Supermix (Bio-Rad) and the following primer sets, Cyclin A2 Fwd 5’- TGAGTTTGATAGATGCTGACCCG-3’, Rev 5-ATCCAGTCTGTTGTGCCAATGAC-3’; Cyclin E2 Fwd 5’-TCTGTGCATTCTAGCATCGACTC-3’, Rev 5’- AAGGCACCATCGTCTACACATTC-3’; E2F1 Fwd 5’-TGCCAAGAAGTCCAAGAATCA-3’, Rev 5’-CTTCAAGCCGCTTACCAATC-3’; Apaf1 Fwd 5’-GATGTGGAGGTGATCGTGAAG- 3’, Rev 5’-GTAGTGTCGTGGTAGGTCAT-3’; Puma Fwd 5’-ATGGCGGACGACCTCAAC-3’, Rev 5’-AGTCCCATGAAGAGATTGTACATGAC-3’; Noxa Fwd 5’- AGGAAGGAAGTTCCGCCG-3’, Rev 5’-AGCGTTTCTCTCATCACATCACA-3’. Reactions were processed in a CFX96 Touch Real-Time PCR System (Bio-Rad). Fold change was determined using the ΔΔ CT quantification method, where Hepatoma-Derived Growth Factor (HDGF) was used as the housekeeping gene.

## Acknowledgments

We thank Dr. Wade Sigurdson, director of the Confocal Microscopy Core at the University at Buffalo, for providing access to and training on the Leica TCS SP8 during quarantine. We also thank Drs. Sarah X. Zhang and Joshua J Wang for generous use of their cryostat. This work was supported by a research grant from the National Institutes of Health to C.S.T. [GM112721].

## Author Contributions

J.C and C.S.T. designed the study. J.C. performed the majority of the experiments and analyzed the data. He was aided in the behavioral studies by B.R. and in the immunofluorescence studies by L.N.M. Z.Y supervised the behavioral studies, and M.L.F supervised and help design the immunofluorescence studies. J.C. and C.S.T. wrote the manuscript with feedback from all authors.

---

**Supplemental Figure 1.**
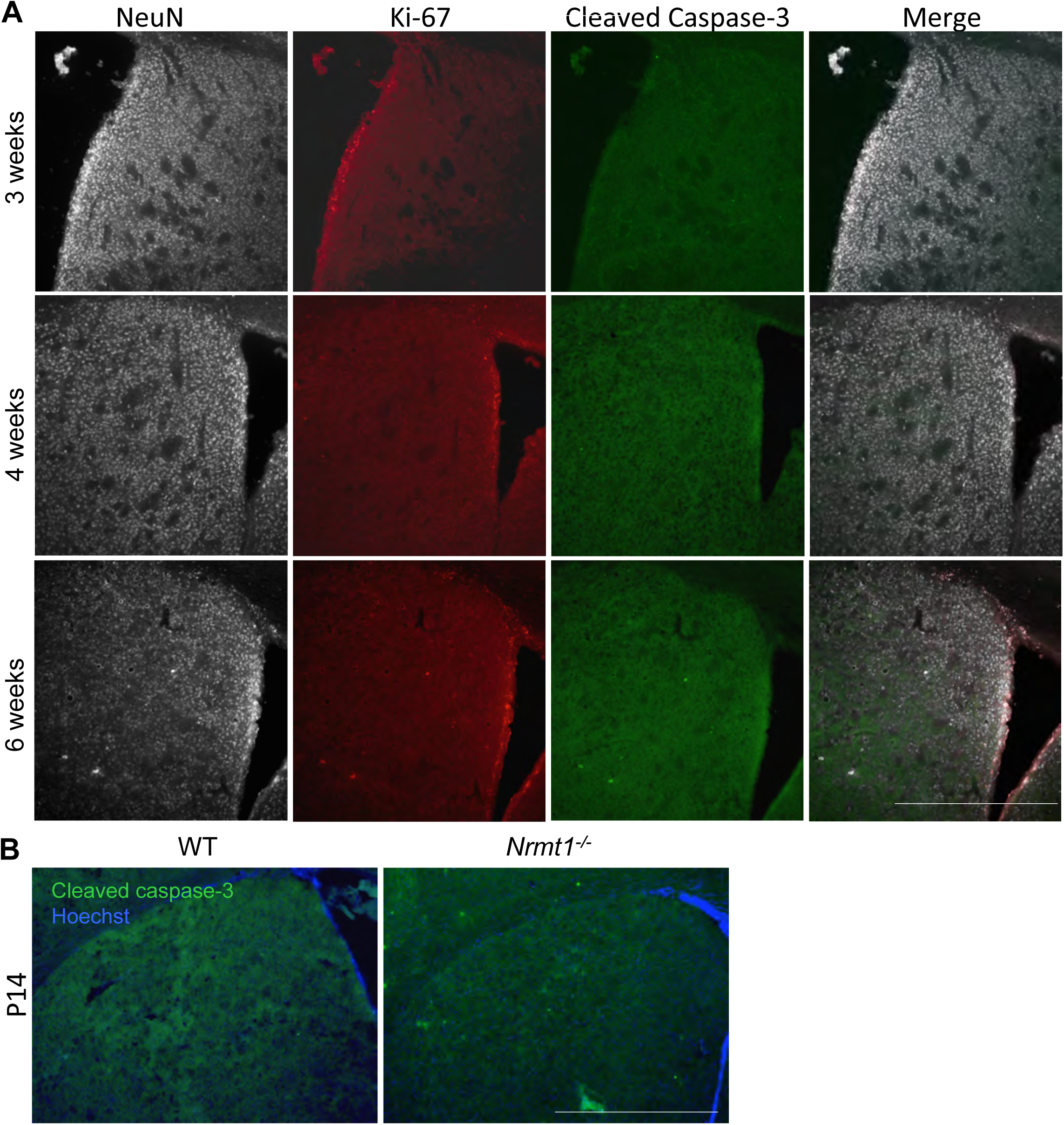
(*A*) WT ventricle/striatal sections triple-stained for NeuN (gray), Ki-67 (red), and cleaved caspase-3 (green). (*B)* WT and *Nrmt1^-/-^* ventricle/striatal sections from P14 mice stained for cleaved caspase-3 (green). Blue is Hoechst counterstain. Scale bar = 1000 µm.

